# SegFinder: an automated tool for identifying RNA virus genome segments through co-occurrence in multiple sequenced samples

**DOI:** 10.1101/2024.08.19.608591

**Authors:** Xue Liu, Jianbin Kong, Yongtao Shan, Ziyue Yang, Jiafan Miao, Yuanfei Pan, Tianyang Luo, Zhiyuan Shi, Yingmei Wang, Qinyu Gou, Chunhui Yang, Chunmei Li, Shaochuan Li, Xu Zhang, Yanni Sun, Edward C. Holmes, Deyin Guo, Mang Shi

**Author notes:** These authors contributed equally to this work (Xue Liu, Jianbin Kong).

## Abstract

Metagenomic sequencing has expanded the RNA virosphere, but many identified viral genomes remain incomplete, especially for segmented viruses. Traditional methods relying on sequence homology struggle to identify highly divergent segments and group them confidently within a single virus species. To address this, we developed a new bioinformatic tool – SegFinder – that identifies virus genome segments based on their common co-occurrence at similar abundance within segmented viruses. SegFinder successfully re-discovered all segments from a test data set of individual mosquito transcriptomes, which was also used to establish parameter thresholds for reliable segment identification. Using these optimal parameters, we applied SegFinder to 858 libraries from eight metagenomic sequencing projects, including vertebrates, invertebrates, plants, and environmental samples. Furthermore, we identified 108 (excluding RdRP) unique viral genome segments, of which 55 were novel and 32 showed no recognizable sequence homology to known sequences but which were verified by the presence of conserved sequences at the genome termini. SegFinder is also able to identify segmented genome structures in viruses previously considered to be predominantly unsegmented, and in doing so expanded the number of known families and orders of segmented RNA viruses, making it a valuable tool in an era of large-scale parallel sequencing.

## Introduction

Segmented RNA viruses maintain their genetic information across multiple distinct RNA molecules and are widespread across diverse hosts including animals, plants, bacteria, and fungi. According to the International Committee on the Taxonomy of Viruses (ICTV), there are over 40 families or genera of segmented RNA viruses, that include a number of notable human, animal, and plant pathogens (https://ictv.global/report/genome). These viruses typically have between 2 and 12 genome segments. For example, members of *Partitiviridae* typically have 2 segments, that of *Orthomyxoviridae* have 6-8, while species within the *Reoviridae* possess 11-12 segments ^1–3^. The evolution of segmentation in these viruses is complex, with the number of segments even varying within the same virus family ^4,5^. These segments are also interdependent, each encoding proteins that fulfill essential roles in the viral life cycle. Consequently, segments typically co-occur at levels proportional to each other ^5^. In addition, although segments encode different proteins, they often possess highly conserved reverse complementary termini or similar regulatory sequences in their non-coding regions ^6^, providing evidence that they belong to the same virus.

Given that the genomic content of segmented viruses is spread among different segments, it is crucial to identify all relevant segments within a virus to assemble a complete genome. Traditionally, individual genome segments are discovered and validated using a variety of molecular or biochemical tools after isolating and identifying virus particles ^1,7–9^. A notable example is reovirus type 3 (i.e., mammalian orthoreovirus 3), in which the first three segments (S, L, M) were identified in a mixture of double-stranded fragments based on several criteria: a distinct melting profile with a specific melting temperature (Tm), resistance to ribonuclease enzymes, sedimentation behavior independent of ionic concentration, and a balanced base composition of A-to-U and G-to-C ^10^. Further research using polyacrylamide gel electrophoresis expanded the reovirus type 3 genome to ten segments, including three large, three intermediate, and four small fragments ^11^. While these methods are effective, they depend on the isolation and purification of virions, which can be challenging for viruses discovered from metagenomic data.

Advancements in high-throughput sequencing and meta-transcriptomics hold great potential for discovering viral genome segments. These methods involve identifying all relevant viral nucleotide or protein sequences from the same sample and determining their single-virus origin based on taxonomic annotations. However, this homology-based approach also faces challenges: it is primarily effective for viruses closely related to known references and struggles to distinguish between segments from related viruses present in the same sample. For divergent viruses lacking clear homology, alternative strategies are required. For instance, segmented viruses can be identified by the similar abundance of each segment and confirmed through the analysis of highly conserved and complementary 5′- and 3′-terminal nucleotide sequences ^6,12–15^. A recent example is our recent work with the discovery of the Jingmen tick virus (JMTV); this virus was identified through similar abundance levels across segments and confirmed via genomic characterization, proteomics, and phylogenetic analyses ^5^.

Abundance-based methods are particularly useful for discovering viral genome segments if multiple sequencing runs are performed ^13–15^. In a recent meta-transcriptomics study that sequenced the viromes of 161 mosquito individuals, Batson and colleagues identified complete genome sets of 27 highly prevalent segmented viruses by co-occurrence and matching abundance levels, including those lacking known protein homology ^16^. Of note, they discovered and validated genome segments 7 and 8 of Wuhan mosquito virus 6 and revealed that Culex narnavirus 1 is a bi-segmented virus. These findings provide important insights into the genomic diversity and composition of insect-specific viruses, about which little was previously known ^16^. While this approach to segment discovery is impressive, it remains labor-intensive, and currently no computational tools or pipelines exist to systematically discover new viruses and their corresponding segments from multiple sequencing data.

RNA virus discovery has surged in recent years, with a major expansion from their known diversity ^14,15,17,18^. However, few of these viruses can be isolated for genome segment identification using *in vitro* methods ^19^. Typically, only the genome segment containing the RNA-dependent RNA polymerase (RdRP) protein is identified, with segments encoding other viral proteins undetected ^18^. Furthermore, due to the complex bioinformatic procedures required to reconstruct the full genome, the search for matching segments is often restricted to a few of selected viruses deemed critical for specific studies. To address this challenge, we developed SegFinder, an automated program specifically designed to identify RNA virus genome segments from parallel sequencing data by analyzing their co-occurrence and matching abundance levels. This method not only detects different segments of known segmented viruses, but also identifies segments encoding previously uncharacterized proteins, referred to as viral “dark matter”. Using SegFinder, we greatly expanded the diversity of viral families and orders of segmented RNA viruses, making this tool valuable in an era of large-scale parallel sequencing.

## Results

### Overview of SegFinder

We developed an automated tool, SegFinder (Segmented Virus Automation Finder), for discovering RNA viruses and their associated genome segments from multiple parallel sequencing data sets. This new tool operates on the principle that all segments from a single virus must co-occur and should exhibit similar abundance levels in each sample (i.e., co-occurrence). Hence, the program identifies complete sets of RNA virus genomes by first identifying the RNA-dependent RNA polymerase (RdRP) segment and then using co-occurrence data to detect the remaining contigs. The first step is the discovery and refinement of segments containing the RdRP gene, including the removal of misassembled regions and screening for the length and abundance of the open reading frame (ORF) to identify true RNA virus sequences for downstream analysis (Figure 1a). Next, SegFinder estimates correlations between each RdRP segment and the remaining contigs by mapping all reads from multiple libraries to the assembled contigs, including the curated RdRP gene segment data sets. This process results in a correlation matrix for each pair of contigs, with correlated segments forming clusters that represent potential segments associated with a single virus species (Figure 1b). The final step in confirmation of the viral segments involves removing contigs containing cellular proteins, as well as those failing to meet length and abundance thresholds. In addition, we excluded clusters that only contained the RdRP segment, those with more than one RdRP segment, and those in which the RdRP segment appeared in fewer than three sequencing libraries (Figure 1c).

**Figure 1.**
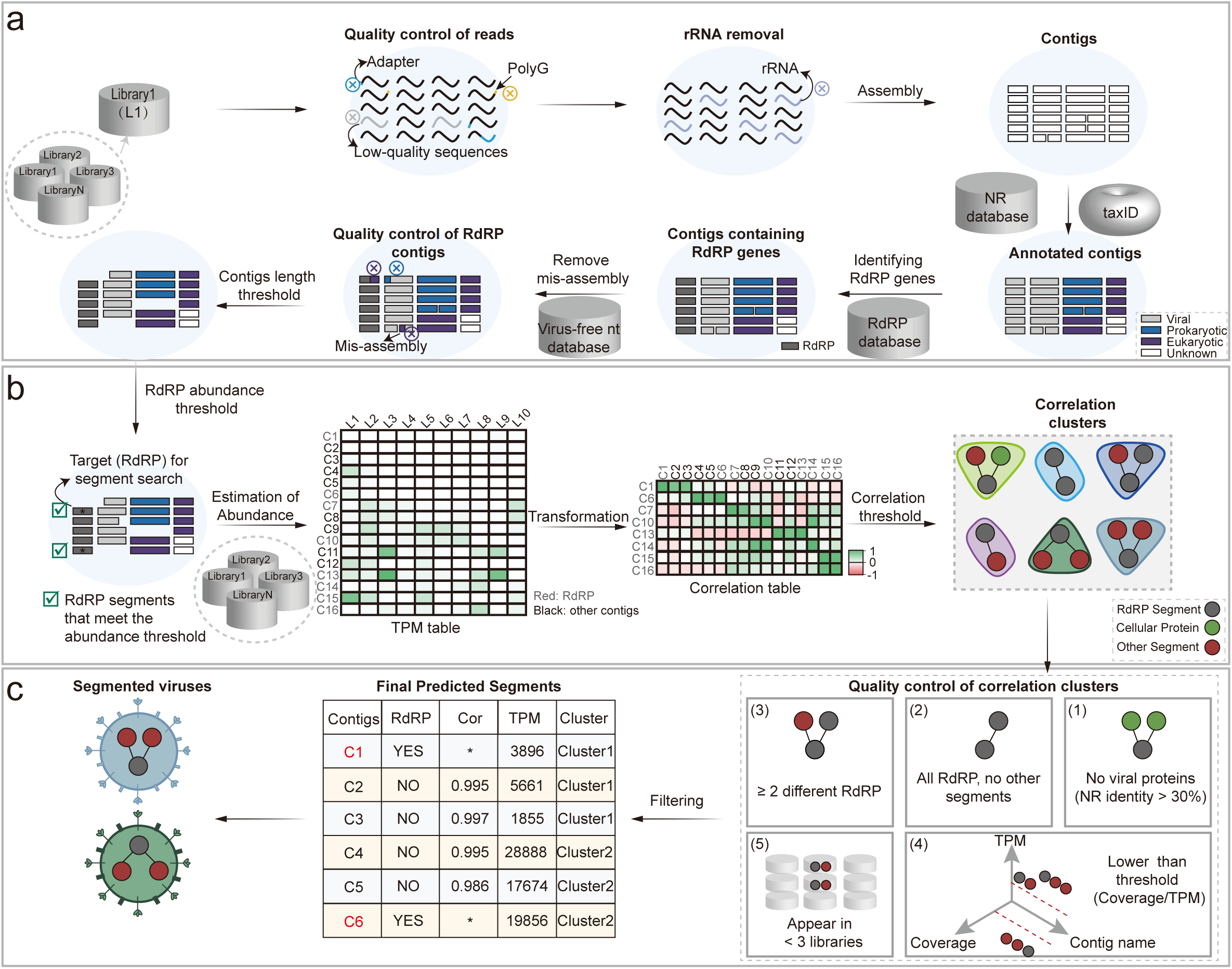
SegFinder workflow. a. Schematic overview of RdRP identification in RNA viruses. The inputs are FASTQ files from multiple meta-transcriptomic libraries. Abbreviations: rRNA (ribosomal RNA), NR (Non-Redundant Protein Sequence Database), NT (Nucleotide Sequence Database). b. The processing pipeline for correlation calculations. Abbreviations: L (Library), C (Contig). c. Schematic illustration of filtering segmented RNA virus clusters. Abbreviations: Cor (Correlation), TPM (Transcripts Per Kilobase Million).

### Optimizing parameter sets to identify viral genome segments

To optimize parameter combinations for RNA virus segmentation detection, we employed SegFinder to a meta-transcriptome data set from mosquitoes in which RNA virus segments are well characterized ^16^. We considered two parameters: the correlation coefficient, measured using Spearman’s test, and the sequencing coverage of the RdRP gene, estimated using the megahit assembly program. Applying various thresholds to RdRP coverage, we observed that false discoveries dropped to zero when coverage exceeded 50 (Figure 2a, Supplementary Figure 1). Setting the coverage threshold at 50, we then analyzed the false negative rates for each virus at different correlation coefficients. This revealed that the last segment of Wuhan Mosquito Virus 6 could not be detected once the correlation coefficient was set at 0.83 (Figure 2b). Consequently, for subsequent analyses, we adopted a coverage threshold of 50 and a correlation coefficient of 0.8 as the default parameter settings.

**Figure 2.**
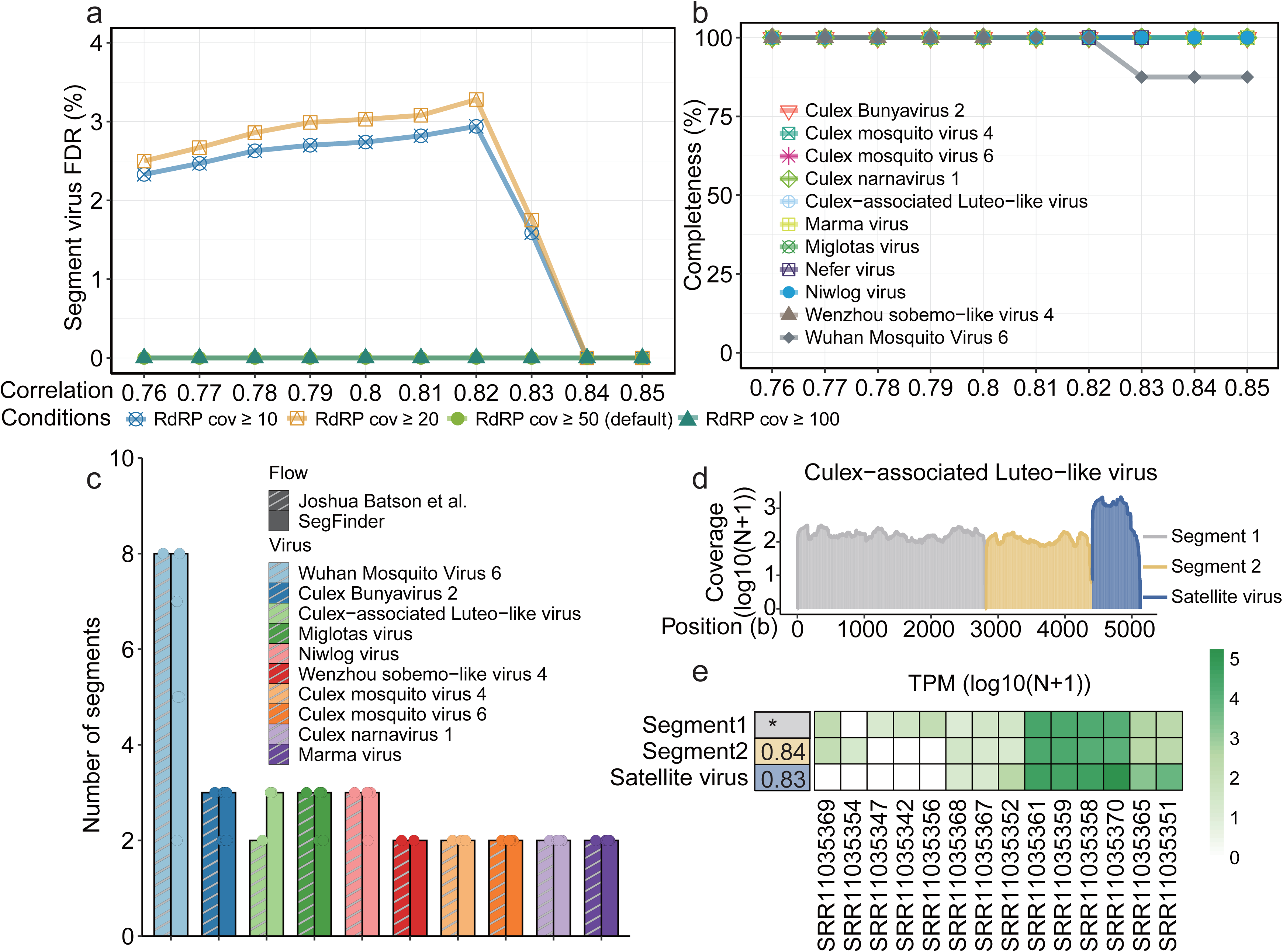
Application of SegFinder to individual mosquito metatranscriptomes to optimize parameter settings. a. False discovery rate for different combinations of SegFinder’s coverage parameters compared to known segmented viruses (cov, coverage). b. Line plots showing the completeness of segmented viruses against varying correlation thresholds. c. Results from the application of SegFinder using the optimal parameters settings: length threshold of 600 bp, TPM threshold of 200, coverage threshold of 50 for RdRP (10 for other segments), and correlation setting of 0.8. The findings from Batson et al. ^16^ are shown by slashed squares, while those from SegFinder are indicated by squares. Dots represent the number of segments found in different libraries. d. Coverage of three genome segments of the newly identified Culex-associated Luteo-like virus. e. Heatmap showing the prevalence and abundance level of Culex-associated Luteo-like virus in each sample.

Our detection pipeline, using the default parameters described earlier, identified ten potentially complete segmented viruses. These included one eight-segmented virus (i.e., Wuhan mosquito virus 6), four three-segmented viruses (i.e., Culex Bunyavirus 2, Culex-associated Luteo-like virus, Miglotas virus, Niwlog virus), and five two-segmented viruses (i.e., Wenzhou Sobemo-like virus 4, Culex mosquito virus 4, Culex mosquito virus 6, Culex narnavirus 1, and Marma virus) (Figure 2c) (Supplementary Table 1). For these viruses, the number of segments identified matched those reported in the original publications, with the exception of Culex-associated Luteo-like virus, for which an additional segment was found (Figure 2c). This segment displayed a similar abundance level and a high correlation coefficient of 0.83 compared to the segment containing the RdRP (Figure 2d-e). However, subsequent analysis based on Gene3D evidence showed that this newly discovered segment was in fact the coat protein of the satellite virus (CATH Superfamily 2.60.120.220). This newly discovered satellite virus occurred at a frequency of 5.59% (9 out of 161) in the library and is 701 bp in length. (Figure 2e).

### Identifying viral genome segments in previously published data

We next used SegFinder to analyze 858 meta-transcriptomes from eight previously published studies, covering a variety of sample types. These included two vertebrate-related (i.e., bats and pangolins), two invertebrate-related (mosquitoes and honeybees), three plant-related (beet, seasonal root, and wheat rhizosphere soil/root), and one environmental (peat) data sets (Figure 3a, Supplementary Table 2). Our analysis revealed 108 (excluding RdRP) unique segments from 45 virus species across 12 orders (Figure 3b, Supplementary Table 3). Among these, 55 segments were novel, and 32 segments showed no recognizable homology to known sequences, which we referred to as “un-annotated” segments. These un-annotated segments were most prevalent in environmental samples (55.56%), followed by plant (31.58%) and invertebrate samples (17.07%), and were least frequent in vertebrate samples (4.17%) (Figure 3c).

**Figure 3.**
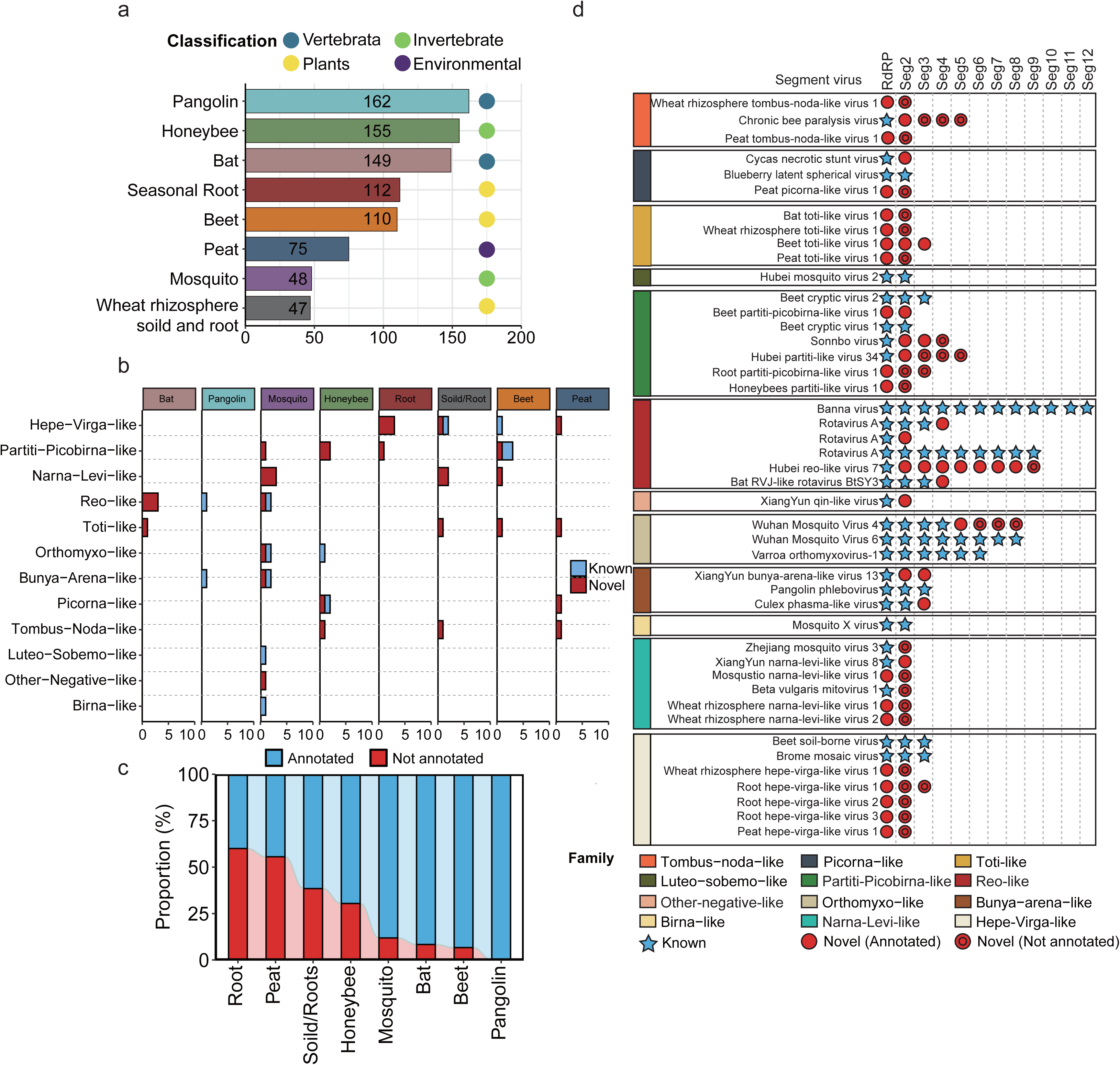
Application of SegFinder to vertebrate, invertebrate, plant, and environmental metatranscriptomic data sets. a. Description of the categories and number of data sets. b. Number of segmented viruses found in all data sets. Blue indicates known segmented viruses; red indicates unknown segmented viruses. c. Percentage of segmented viruses that can be annotated by the nr and nt databases, based on the number of segments. d. Characterization of segmented viruses in various sample types. The colors in the left box represent the superclades to which the segmented viruses belong. Virus segments are represented by blue pentagrams for known segments, red circles for new segments that can be annotated by the nr and nt libraries, and red double circles for new segments that cannot be annotated by the nr and nt libraries.

These results show that SegFinder can identify both known and uncharacterized virus segments in multi-segmented families such as the *Reoviridae* and *Orthomyxoviridae*. For example, it identified all 12 segments of the Banna virus genome within the *Reoviridae*, which has been identified previously using experimental methods. In addition, it identified 8 segments in Wuhan mosquito virus 4, four more than described previously. One of these segments exhibited 32.7% amino acid similarity with hypothetical protein 1 of Byreska virus, while the other three are previously unannotated (Figure 3d). Similarly, Hubei reo-like virus 7, previously only known by its RdRP segment, now has eight newly recognized segments.

More importantly, for several viruses, SegFinder has uncovered additional, highly divergent segments beyond what was recognized as typical number of genome segments (Figure 3d and 4). For instance, Chronic bee paralysis virus (CBPV), traditionally considered to have two segments, was identified to contain three additional unannotated segments of lengths 1040, 982, and 634 bp, respectively. These co-occur with the RdRP segments in over 150 libraries, suggesting that they belong to the same virus. Interestingly, some segments of CBPV are absent in some of the certain libraries. For instance, segments 3 and 4 were missing from two samples in which the other segments were relatively abundant (Supplementary Figure 2). This could be due to recombination, although the possibility that these segments are part of a satellite virus cannot be completely excluded. In addition, we identified six bi-segmented narnaviruses, even though this group was typically thought to possess one genome segment aside from a few notable exceptions^16,20^ (Figure 3d). Furthermore, members of the partiti-like virus group, such as Sonnobo virus and Hubei partiti-like virus 34, now display four and five segments, respectively, exceeding the typical two and occasionally three segments observed within this group (Figure 3d and 4).

**Figure 4.**
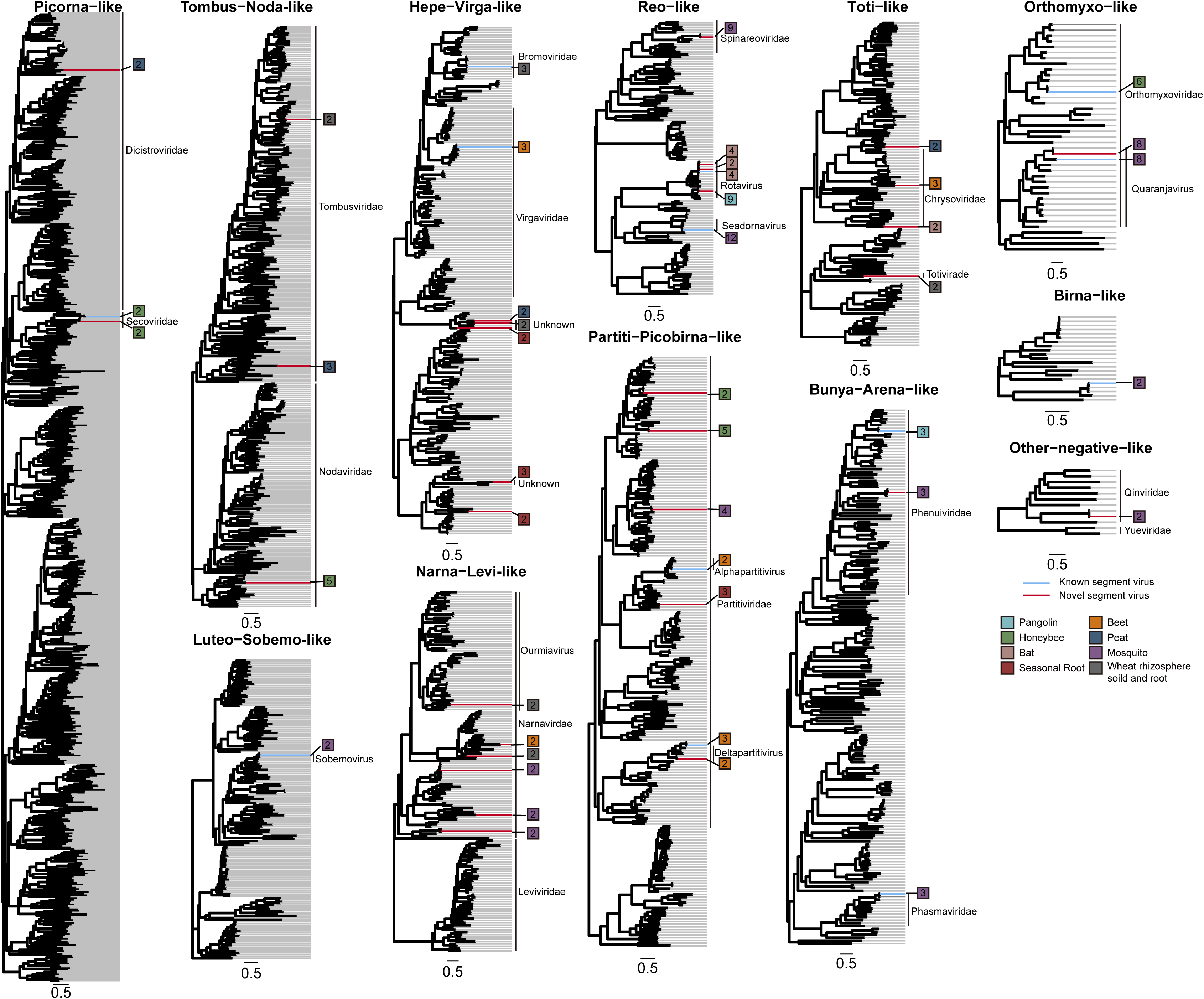
Phylogenetic trees illustrating the evolutionary position and segment numbers of the viruses identified in this study. Phylogenetic trees were estimated at the virus supergroup level using a maximum likelihood method based on analysis of the RdRP protein. The trees are mid-point rooted. Within each tree, existing virus species are marked with blue lines, while newly identified segmented viruses are marked with red lines. The labeling on the right shows the families or genera within each clade, the types of libraries used, and the number of segments found in each segmented virus. Each scale bar represents 0.5 amino acid substitutions per site.

### Verification of viral segments

An additional analysis was performed on each identified viral segment to confirm its association with the same viruses as the RdRP (Figure 5a). Accordingly, all the 108 segments discovered were manually checked for co-occurrence using sequence mapping data (Figure 5b, Supplementary Table 3). We also assessed whether the segments discovered had homology to proteins encoded by related viruses; this accounted for 70.37% (76/108) of all segments (Figure 5a). For cases in which the segments discovered had no homology to proteins from known viruses, we employed a homology-independent approach for confirmation, searching for highly conserved motifs or complementary nucleotide sequences at the 5′- and 3′-termin of the virus genomes. Using the MEME program, we found that 21 of the 45 segmented viruses had conserved motifs at their termini (Figure 5a). For instance, three related members of the *Partitiviridae* —Beet cryptic virus 1, Beet cryptic virus 2, and Beet partiti-picobirna-like 1— exhibited several highly conserved sequence motifs, including a 13 bp stretch conserved across all segments (i.e. Forward: AGATCGGAAGAGC/Reverse: GCTCTTCCGATCT) (Figure 5c). Overall, 84.26% 91/108) of all newly identified segments were confirmed through homology or conserved termini.

**Figure 5.**
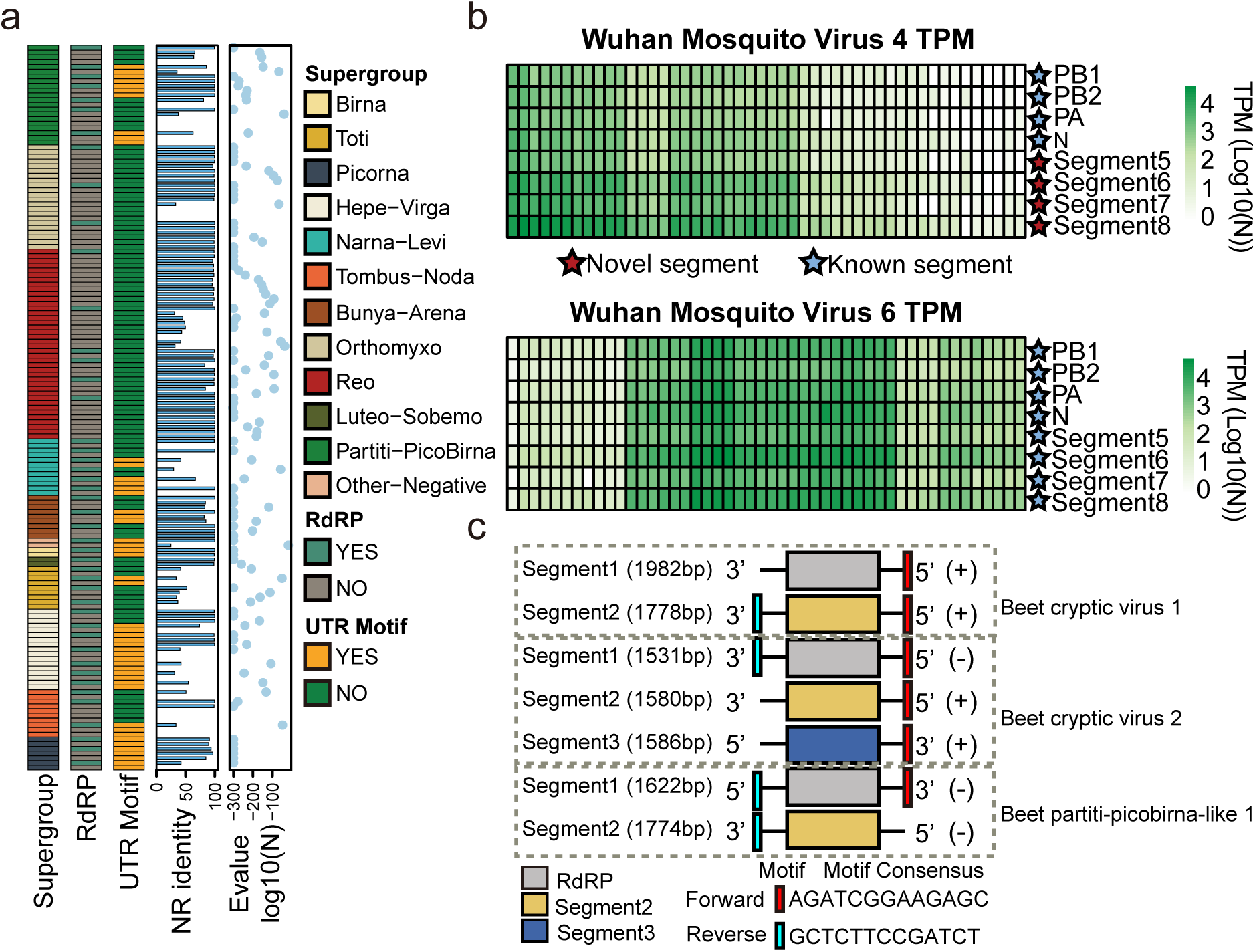
Verification of the viral segments detected. a. The characterization of the segmented viruses and their segments found in the metatranscriptomic libraries. b. TPM values for the eight genome segments of the Wuhan mosquito virus 4 and Wuhan mosquito virus 6 across 48 libraries (represented by vertical green squares). c. Schematic representation of conserved untranslated regions discovered in Beet cryptic virus 1, Beet cryptic virus 2, and Beet partiti-picobirna-like 1.

## Discussion

A number of powerful software solutions exist for RNA virus detection in the metagenomic era, including VirFinder, DeepVirFinder, VirSorter, and VirSorter2 ^21–24^. While these tools are effective for identifying RNA virus genomes or genome segments, they often struggle to accurately assign the segments identified to same virus species or detect highly divergent genome segments that lack sequence homology to known proteins. In contrast, our program – SegFinder – is able to identify both the RdRP segment and the corresponding genome segments of newly discovered viruses without relying on sequence homology, as commonly used in many studies ^5,13,16^. Indeed, while homology-based approaches are useful for identifying closely related segments, they face several challenges. First, viral genes can evolve at very different rates ^25^. Although RdRP genes are relatively well conserved, other viral proteins, such as the capsid and glycoproteins, evolve more rapidly, leading to extensive sequence divergence even within the same virus family. For instance, while the L proteins of Lassa virus, Boa Av NL B3 virus and salmon pescarenavirus 1 (all within *Arenaviridae*) show clear sequence homology, their glycoproteins and nucleoproteins do not ^15,26,27^. As a result, the discovery of a divergent member of a segmented virus is often challenging. Second, genes within a single virus can exhibit distinct evolutionary histories due to genomic recombination, reassortment, or horizontal gene transfer, as shown by differing phylogenetic histories of the RdRP and structural genes ^14^. In extreme cases, a virus may possess genetic components from both DNA and RNA viral origins ^28^, complicating the association of non-RdRP segments with the RdRP. Accurately capturing the essential features of segmented viruses, particularly their occurrence, is therefore crucial for revealing their complete genomes. Lastly, a homology-based approach may fail to accurately assign segments when two related viruses are present in the same sample with similar abundance levels. However, our method, which relies on patterns of change rather than abundance levels within a single sample, can effectively resolve this issue.

The method used here, based on co-occurrence, requires a data set in which the virus is highly prevalent across many sampled populations (i.e., sequenced libraries) and exhibits varying levels of abundance. In addition, samples with greater abundance should provide sufficient read depth to enable the assembly of complete viral genomes, enhancing the robustness and accuracy of segment discovery. As a consequence, we implemented stringent default parameter settings to minimize false positives, setting high thresholds for correlation (≥ 0.8), prevalence (appearance in ≥ 3 libraries), abundance (RdRP coverage ≥ 50), and completeness of potential viral segment (contig length ≥ 600 bp). A higher correlation threshold helps prevent false positives that might arise from a positive association between genomes of two different organisms rather than within a single virus, as inter-organismal genomic correlations are typically lower than those within a single organism. The prevalence threshold aims to avoid spurious correlations due to insufficient data, while strict abundance and completeness thresholds help prevent the misidentification of fragmented assemblies of unsegmented virus genomes as segmented ones due to low coverage. However, these stringent thresholds may reduce the number of segmented viruses identified. For example, viruses in the *Spinareoviridae*, which typically have 9 to 12 genome segments ranging from 0.5 to 4.8 kb in length, might see their smallest segments excluded due to the 600 bp length threshold. Additionally, setting no abundance threshold enabled the discovery of novel segmented flaviviruses that share 57.34% amino acid similarity to Inopus flavus jingmenvirus 1 in the RdRP region ^29^. Therefore, finding the optimal parameter settings to balance false positives and negatives is challenging, although parameters can be adjusted for more sensitive discovery.

SegFinder is also to identify multipartite RNA viruses. Indeed, we identified members of the *Partitiviridae* (dsRNA) and *Secoviridae* (ssRNA (+)), both of which are multipartite ^30^. In multipartite viruses, each segment exhibits different abundance levels, which can vary by more than an order of magnitude between segments encoding structural proteins and those encoding non-structural proteins like the RdRP ^5,16^. However, such disparities do not impact co-occurrence estimations since the differences are proportional. Additionally, SegFinder offers versatile applications beyond identifying viral genome segments. By adjusting the abundance or completeness thresholds, the program can detect fragmented contigs of a single unsegmented virus genome. This allows users to assemble a complete set of genomic fragments, which can serve as templates for primer design in PCR assays aimed at obtaining complete genomes. Moreover, as demonstrated here, SegFinder can also identify satellite viruses that occasionally co-occur with the virus under investigation. These viruses often encode highly divergent proteins that may be overlooked in typical virus discovery efforts.

SegFinder was also able to reveal the complexity of genome evolution in some groups of RNA viruses and show that segmentation is more commonplace than previously anticipated. Indeed, this and other recent work have greatly expanded the diversity of segmented viruses, as highlighted by the recent discovery of novel segmented viruses related to those with unsegmented genomes, such as the Jingmenviruses (*Flaviviridae*) ^4,5^ and bi-segmented coronaviruses from aquatic vertebrates ^31^ and environmental samples ^32^. Furthermore, we increased the number of segments within groups of known segmented viruses, demonstrating that viral genome organizations have greater flexibility than previously realized. For example, partitiviruses, that are traditionally considered to be bi-segmented, are now demonstrated to have segments numbers ranging from 2 to at least 5.

Despite its utility, SegFinder has several limitations. First, it requires the parallel sequencing of multiple samples, within which the target virus must have a relatively high prevalence and a moderate abundance level. Metagenomic projects and viruses meeting these criteria might be limited in number. Second, our program provides an end-to-end pipeline that includes assembly, mapping, and annotation processes, which demand moderate computational resources and relatively long computation times. Thirdly, the presence of conserved sequences at genome termini cannot verify all detected segments due to potential issues with sequence assembly quality, but a follow-up Rapid Amplification of cDNA Ends (RACE) assay could help address this problem. Finally, while SegFinder can identify new segments with high precision, even when there is no homology to existing proteins, it cannot provide structural or functional information, necessitating additional methods for these analyses. Nonetheless, SegFinder captures the key features of segmented viruses and can reveal complete virus genomes, thereby providing information that is crucial to understanding virus function and evolution.

## Material and methods

### Reference databases

To enhance the accuracy of segmented RNA virus identification, we developed a curated RdRP (RNA-dependent RNA polymerase) database and a nucleotide database devoid of viral sequences. The steps involved in the development of each database are described below.

### RdRP database

The RdRP database utilized in this study was developed through updates to an database developed by our group ^14^. Sequences within this database were re-annotated and filtered using the Pfam A database to retain only the core regions of RdRP ^33^ (e-value 1E-5). This enabled the creation of a refined, high-quality database specifically for identifying RNA viruses.

### Virus-free non-redundant nucleotide (virus-free nt) database

The no-viral NT (i.e., virus-free) database was initially constructed by eliminating sequences identified as viral based on their IDs ^34^. Subsequently, any remaining viral sequences were removed using BLASTn against representative viral genomes from NCBI.

### Metatranscriptomic data sets

Nine publicly available meta-transcriptomics sequencing data sets were downloaded from NCBI SRA database, comprising mosquitoes (PRJNA605178, PRJNA778885), honeybees (PRJNA706851), bats (PRJNA929070), pangolins (PRJNA845961), beet (PRJNA808220), seasonal root (PRJEB35805), wheat rhizosphere soil and roots (PRJNA880647) and peat (PRJNA386568) ^16,35–41^ (Supplementary Table 2). Among these, the mosquito data set (PRJNA605178) was used to establish standard parameters that maximize the discovery rate of new virus segments. The remaining data sets served as diverse examples in which additional genomic segments were identified.

### Quality control steps for reads

Quality control and the preprocessing of raw sequencing reads were performed using fastp (version 0.22.0) and RiboDetector (version 0.2.7), which efficiently remove ribosomal RNA reads ^42,43^. Subsequently, bowtie2 (version 2.2.5, set to -- local) was employed to remove host reads ^44^ if the host genome accession number is provided by the user.

### Identification of the viral RdRP segment

After removal of rRNA reads, the remaining reads were *de novo* assembled using MEGAHIT (versions 1.2.9) or MetaSPAdes (version 3.13.0) ^45,46^, depending on user’s choice. The assembled contigs were then compared against the non-redundant (nr) protein sequence database using DIAMOND (version 0.9.19) ^47^, with an E-value cutoff of 1E-4 to balance high sensitivity with a low false-positive rate. Contigs potentially containing viral genome sequences were identified for further blastx analysis against the RdRP protein database, which identified RdRP-containing contigs. Subsequently, each RdRP-containing contig was compared against the virus-free nt database using the BLASTn program for the identification of chimeric sequences between virus and non-viral sequences, as well as mis-assembled sequences, which were subsequently removed. In the end, only high quality RdRP-containing contigs were retained for further characterization.

### Abundance estimations

We estimated the abundance of the entire set of viruses contigs across all libraries. To streamline this analysis, we applied a length threshold of over 600 nucleotides to the contigs, which enhanced computational efficiency. Subsequently, we utilized cd-hit (version 4.8.1) ^48^ to reduce redundancy among the contigs, setting a threshold of 0.8. In addition, we used ORFfinder (version 0.4.3) with the settings -ml 30 -s 2 to verify that each contig encoded proteins. The relative abundance of each contig was then estimated using the Salmon software (version 0.13.1). This process involved calculating the number of mapped reads per kilobase million reads (TPM) ^49^ for each library, expressed as Transcripts Per Kilobase of per Million mapped reads divided by the transcript length in kilobases, and multiplied by 10^6. Finally, the TPM data were compiled into an abundance matrix for further analysis.

### Correlation analysis

We first removed non-viral portions of the sequence contigs by performing BLASTn analyses against a virus-free nt database. We calculated the Spearman correlation coefficient using the ‘psych’ package in R to analyze the TPM expression matrix. Segmented virus genome clusters were formed based on the correlations (p-value < 0.05) between the viral RdRP gene and other contigs. Specifically, contigs that exhibited a correlation coefficient above the set threshold with the RdRP gene were grouped into a single cluster with the RdRP. These sequences were considered potential segments associated with a single viral species.

### Quality control of the results

We identified and removed the following clusters: (1) clusters primarily comprising non-viral genes that exhibited over 30% amino acid identity with cellular proteins; (2) clusters only containing the RdRP segment; (3) clusters with more than two RdRP segments, indicating the presence of more than two viral species; (4) clusters whose segments fell below the established abundance threshold for comparisons; and (5) clusters where the RdRP segment appeared in fewer than three libraries. The clusters that met our criteria were retained for further analysis and presented in the Results.

### Phylogenetic analysis

We selected a diverse set of representative RdRPs from a large-scale database for phylogenetic analysis ^14,15^. The RdRPs of segmented viruses were aligned using the L-INS-I algorithm in Mafft v7.520 ^50^. The aligned sequences were further processed using the Trimal software^51^ to remove ambiguously aligned regions. We then performed phylogenetic analyses using the maximum likelihood (ML) approach implemented in IQ-TREE v2.2.3 ^51,52^, utilizing the optimal amino acid substitution model. Additionally, we utilized ‘ggtree’ and ‘cowplot’ to root the tree, and for the visualization and arrangement of the tree figure ^53^.

### Verification of segmentation

To confirm that the newly discovered segments belong to the same virus, we initially mapped the sequencing reads to the complete genome set to reassess abundance levels and verify co-occurrence. We then translated and annotated these segments using the blastx program to determine homology and associated functions. Additionally, we identified conserved untranslated regions (UTRs) in each segment, providing further evidence of their linkage to the same virus ^54^.

## Data availability

The segmented virus genomes identified in this study can be accessed through the Figshare link https://figshare.com/articles/dataset/segmented_RNA_virus/26531344.

## Code availability

The original source code for SegFinder is stored at GitHub and figshare repository (https://github.com/Kongloner/SegFinder; https://figshare.com/articles/software/_b_SegFinder_software_code_b_/26770885?file=48632488).

## Acknowledgements

This study is funded by National Natural Science Foundation of China (82341118, 32270160), Natural Science Foundation of Guangdong Province of China (2022A1515011854), Shenzhen Science and Technology Program (JCYJ20210324124414040, KQTD20200820145822023), Hong Kong Innovation and Technology Fund (ITF) (MRP/071/20X), Major Project of Guangzhou National Laboratory (GZNL2023A01001 & GZNL2023A01008), Guangdong Province “Pearl River Talent Plan” Innovation, Entrepreneurship Team Project (2019ZT08Y464), and the Fund of Shenzhen Key Laboratory (ZDSYS20220606100803007). E.C. Holmes was supported by an NHMRC (Australia) Investigator Award (GNT2017197) and by AIR@InnoHK administered by the Innovation and Technology Commission, Hong Kong Special Administrative Region, China.

## Author contributions

Conceptualization, X.L., E.C.H., D.-Y.G., and M.S.; Methodology, X.L., J.-B.K., Y.-F.P., E.C.H., D.-Y.G., and M.S.; Software compilation, X.L., J.-B.K., Z.-Y.Y., Y.-T.S., J.-F.M., S.-C.L., X.Z., Y.-N.S, E.C.H., D.-Y.G., and M.S.; Writing – original draft, X.L. and M.S.; Writing – review and editing, All authors; Funding acquisition, D.-Y.G., and M.S.; Resources (data sets), X.L., J.-B.K., Z.-Y.Y., Y.-T.S., and J.-F.M.; Resources (computational), S.-C.L., X.Z., D.-Y.G., and M.S.; Supervision, Y.-N.S, E.C.H., D.-Y.G., and M.S..

## Supplementary material

**Supplementary Figure 1.**
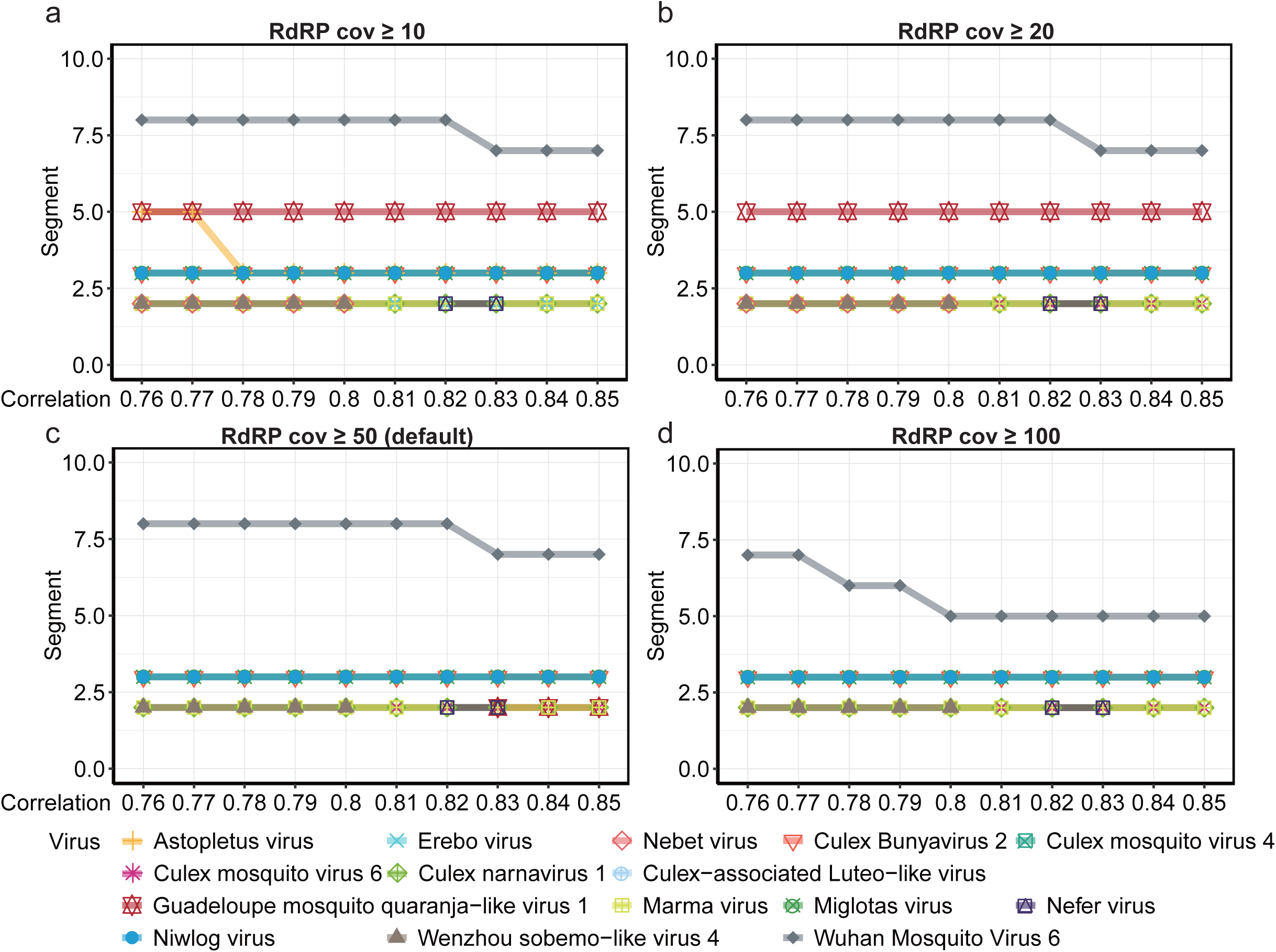
Impact of sequencing coverage on the detection of virus segments in the mosquito metatranscriptome data. The number of identified segmented viruses and their segments at RdRP coverage levels of (a) 10, (b) 20, (c) 50, and (d) 100.

**Supplementary Figure 2.**
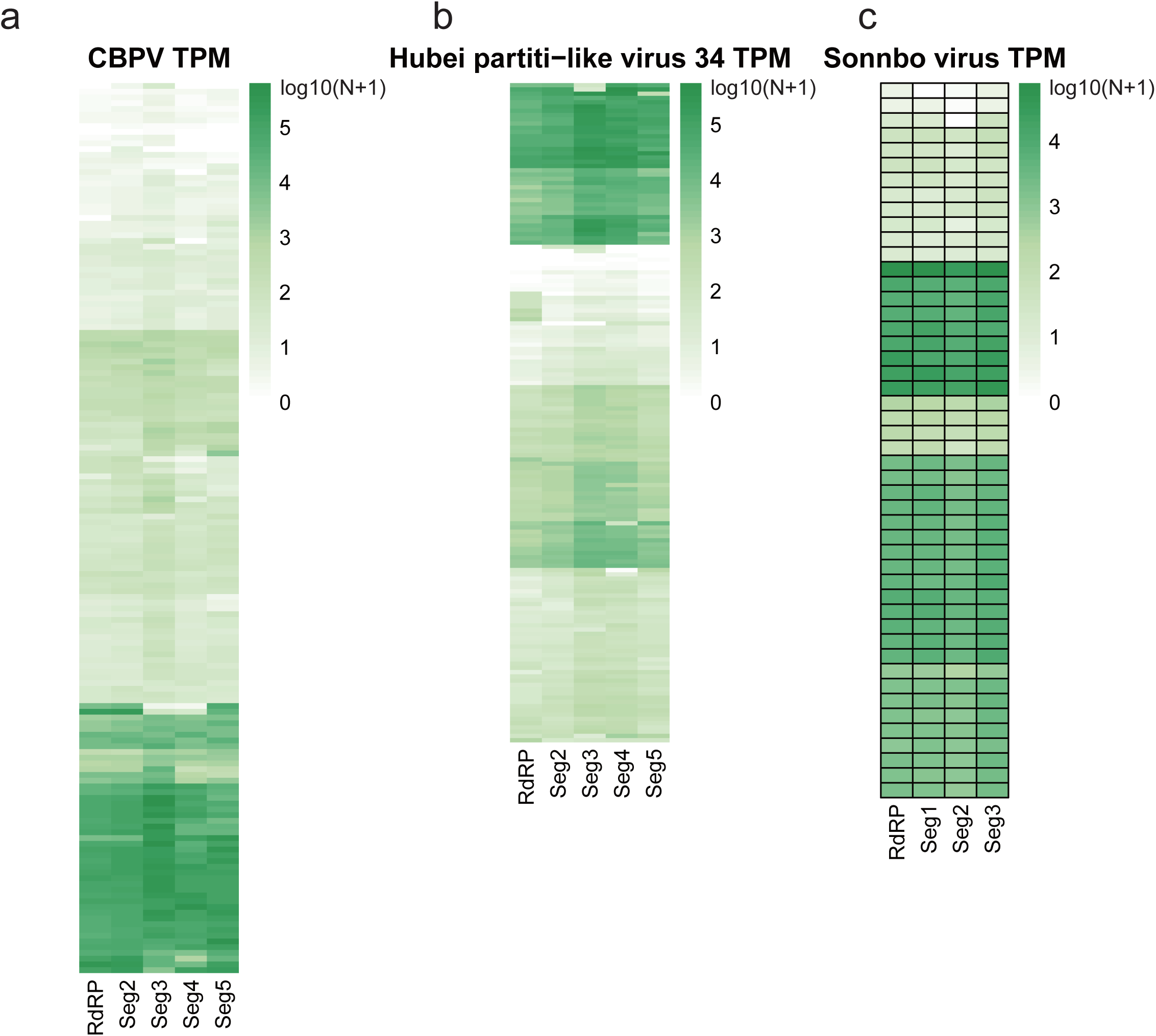
TPM validation of segmented viruses of interest. lAbundance distribution of (a) chronic bee paralysis virus (CBPV), (b) Hubei partiti-like virus 34, and (c) Sonnbo virus segments in their respective data sets.

**Supplementary Table 1. Segmented viruses identified in the individually sequenced mosquito transcriptomes (PRJNA605178)**

**Supplementary Table 2. Data sets used in this study**

**Supplementary Table 3. Detailed information on the virus segment identified from previously published data**

